# Temporal and cell-specific changes to cellular iron sequestration and lipid peroxidation in a murine model of neonatal hypoxic-ischemic brain injury

**DOI:** 10.1101/2025.07.02.662653

**Authors:** Joseph Vithayathil, Anjali Shankar, Ginger L Milne, Frances E Jensen, Delia M Talos, Joshua L Dunaief

**Author notes:** **Corresponding author**: Joseph Vithayathil MD PhD, Division of Neurology, Children’s Hospital of Philadelphia 3500 Civic Center Drive, Philadelphia, PA 19104. Equal contribution (Delia M Talos, Joshua L Dunaief). **Disclaimers:** Views expressed in the article are those of the authors and not an official position of their institutions or funders. **Role of funding source:** Funding sources provided resources and reagents for all conducting all experiments and data analysis noted in this manuscript.

## Abstract

**Background:** Iron accumulation and lipid peroxidation are pathophysiologic mechanisms that drive neonatal hypoxic-ischemic (HI) brain injury. Characterization of spatiotemporal changes in these processes will help elucidate their role in ischemic neuronal injury as an initial step towards developing targeted interventions.

**Methods:** HI was induced in post-natal day 9 mice using the modified-Vannucci model. Hippocampal tissue from ipsilateral HI exposed, contralateral hypoxia exposed and sham animals was collected at 6h, 24h, 72h and 7d post-HI. Tissue was subsequently evaluated for markers of cell death (TUNEL), intracellular iron changes (FerroOrange, fluorescent *in situ* and immunofluorescence), and lipid peroxidation (real time PCR, Gpx4 immunofluorescence and mass spectrometry). Mass spectrometry measured isoprostanes (15-F_2t_-IsoP) and neuroprostanes (4-F_4t_-NP) as lipid peroxidation markers of arachidonic (ARA) and docosahexaenoic acid (DHA), respectively.

**Results:** Compared to sham, the HI hippocampus showed increased intracellular labile iron levels that was maximal at 6h post-HI with subsequent elevation in only neuroprostanes at 24h post-HI. TUNEL labeling peaked at 24h post-HI. At 72h, labile iron levels and lipid peroxidation declined corresponding with peak infiltration of ferritin positive microglia/macrophages and the start of TUNEL staining decline. In addition, surviving neurons had increased expression of Gpx4 peaking at 72h post-HI that normalized by 7d post-HI.

**Conclusions:** These findings suggest that following HI, an acute increase in labile iron and DHA peroxidation are correlated with markers of cell death that peak at 24h post-HI. Microglial/macrophage iron sequestration and neuronal antioxidant responses may ameliorate further injury and represent targets for neuroprotective therapies.

## Introduction

Neonatal hypoxic-ischemic (HI) brain injury affects 1.5 children per 1000 live births in the United States and 3/1000 live births in developing countries resulting in long-term neurodevelopmental disabilities.^1,2^ To date, the only protective therapy that has been developed is therapeutic hypothermia, which is only effective in children with moderate to severe injury and has limited efficacy in low/middle-income countries.^3–5^ The majority of current clinical trials are based on incremental changes to therapeutic hypothermia. Thus, there is a need for development of additional neuroprotective treatments that not only supplement the effects of hypothermia, but also provide benefit to populations where hypothermia is ineffective. In order to begin to develop therapeutic strategies, a better understanding of the pathophysiologic mechanisms of injury is necessary.

Iron dysregulation and lipid peroxidation have been strongly implicated in neonatal HI brain injury as prior work has shown iron accumulation in the areas of ischemic injury as early as 4 hours post insult.^14,15^ The accumulation of iron appears to have toxic effects as treating mice with iron chelators immediately after injury induction ameliorates injury severity.^16,17^ The presumed mechanism is that free iron in the labile iron pool (LIP) reacts with hydrogen peroxide to form hydroxyl radicals, which are the most toxic of all free radicals, in the Fenton reaction.^9–11^ One of the targets of hydroxyl radicals are polyunsaturated fatty acids (PUFAs) which can lead to ferroptosis, a form of programmed necrosis that has recently been observed in neonatal brain injury.^6,7^ It is the result of iron induced peroxidation of polyunsaturated fatty acids resulting in destruction of mitochondrial and cellular membranes.^7,8^ However, the toxic effects of the Fenton reaction and lipid peroxidation can also promote oxidative stress induced injury that results in other forms of cell death including apoptosis.^12,13^ Specific peroxidase enzymes such as glutathione peroxidase 4 (GPX4) are able to protect against cell death by neutralizing lipid peroxides, which can be re-incorporated into cellular membranes. Thus, iron toxicity and lipid peroxidation can contribute to multiple forms of cell death and cell injury.

Despite the strong evidence that these molecular mechanisms contribute to neonatal HI, there is still little understanding of the temporal changes to the pathways regulating iron metabolism and lipid peroxidation following neonatal HI. Specifically, identification of labile iron changes and their correlation with markers lipid peroxidation and neuronal cell death. In addition, the potential cell specific mechanisms involved remain unclear, as different cell types in the developing brain may have selective vulnerability to lipid peroxidation and may differentially regulate iron metabolism. Thus, a better understanding of these cell specific and temporal changes will help determine a therapeutic window for targeting these pathologic mechanisms while also identifying cell-specific targets for future therapeutic development.

The goal of this study is to identify how labile iron pools change and the cell-specific alterations to iron metabolism, while also correlating them with changes in lipid peroxidation and Gpx4. Correlating the changes in these markers is helpful to identify how these responses relate to the time-course of cell death following neonatal HI to help develop future strategies for therapeutic intervention.

In this study, we used an established neonatal HI mouse model to show that labile iron pools and lipid peroxidation are early pathologic mechanisms that occur within the initial 24h after injury, but decline by 72h post-HI. The high labile iron and lipid peroxidation species precede the peak levels of cell death in the hippocampus and the decline in labile iron and lipid peroxidation correlates with an increase in iron sequestration by microglia/macrophages and an increase in neuronal Gpx4 immunoreactivity. These data suggest that endogenous compensatory mechanisms aimed at limiting the extent of brain damage could be augmented or accelerated by therapeutics as a means of acute or preventative neuroprotection.

## Methods

### Animals

All mice were on C57/Bl6N background and were obtained from Charles River Laboratory. Animals were housed in vivarium on 12h light/dark cycle with ad libitum access to food and water. Both male and female mice were included in equal numbers. All mouse litters utilized were derived from in-house breeding of mice. All protocols and treatment were approved by IACUC committee.

### Hypoxia-ischemia procedure

Hypoxic ischemic injury was induced on post-natal day 9 (PND9) mice as previously described.^18,19^ In brief, neonatal mice were separated from dam and had topical lidocaine administered to ventral neck. Mice were then anesthetized using 3.5% isoflurane with maintenance of 1.5%. During surgical procedure, mice were placed on a heating pad to maintain their body temperature. Ventral neck was sterilized with propidium iodide and 70% ethanol prior to skin incision (3-4mm). Forceps were used to expose the right carotid sheath, and the right common carotid artery was isolated with care to avoid damaging of the sympathetic ganglia and vagus nerve. Once exposed, two 5-0 vicryl sutures were used to tie off the carotid artery. Neck incision was then closed with tissue glue and the mouse was placed in a recovery chamber on a heated pad. After a 60-minutes recovery, mice were placed in a hypoxia chamber on heating pad. A Biospherix Oxygen controller was used to fill the chamber with nitrogen and decrease oxygen levels to 8% over 5 minutes. Animals remained in 8% oxygen for 40 minutes and then allowed to recover for 10 minutes on a heating pad before returning to dams. Sham animals underwent exposure of carotid artery under anesthesia, but no ligation of carotid artery and remained in a recovery chamber with no hypoxia exposure. The body temperature of all animals was maintained at 36-38°C throughout the experiment . Animals were then euthanized by decapitation at 6 hours (6h), 24 hours (24h), 72 hours (72h) and 7 days (7d) post-injury.

### Immunofluorescence and immunohistochemistry

Following euthanasia of mice, brains were dissected out and drop fixed in 4% paraformaldehyde for 24h at 4°C. Brains were then transferred through sucrose gradient into 30% sucrose. Tissue then embedded in Optimal Cutting Temperature (OCT) and cryosections generated on Leica 3050S cryostat. 20um sections were mounted onto super frost slides. Immunofluorescence was performed as previously described, but sections were rehydrated in PBS and then underwent antigen retrieval at 95C in AR solution (10mM Citrate Buffer with 0.1% Tween20, pH6). Sections then blocked (5% NDS, 1% BSA, 0.2% Triton-X100 in PBS) for 90minutes at room temperature. Sections then stained with primary antibody (see Supplementary Table 1) diluted in blocking solution and incubated at 4C overnight. Alexa Fluor donkey secondary antibodies used and diluted in blocking solution 1:500 and incubated at room temperature for 2 hours. For Gpx4 staining, Akoya Opal tyramide fluorescent signal amplification kit (NEL840001KT) was used in order to improve antibody signal. Slides then washed and mounted using Fluoromount with DAPI. For all cell counts and quantification, 20x images were obtained from caudal and rostral hippocampus separated by 800um and averaged together. Image analysis was completed using Qupath^20^ for cell counts and mean fluorescent intensity analysis.

### Fluorescent in situ hybridization

For *in situ* staining, ACDbio RNAscope assay was used with Ftl1 (Cat. 314771) and Tfrc (Cat. 427931-C2) probes. Standard RNAscope protocol was used for this assay using Multiplex v2 reagents (Cat. 323100). Images were analyzed using Qupath software, where DAPI positive cells within CA1 pyramidal region were identified and average mean fluorescent intensity of probe signals were measured in each cell.

### Terminal deoxynucleotidyl transferase dUTP nick end labeling (TUNEL)

For TUNEL staining, slides with mounted sections were treated using ThermoFisher Invitrogen kit (C10617). In brief, slides washed in phosphate buffered saline (PBS), baked at 40°C for 20minutes and then post-fixed in 4% paraformaldehyde (PFA) in PBS for 15minutes. Slides then washed in PBS twice. Sections then incubated in proteinase K, followed by incubation in EdU and then washed and incubated with Alexa Fluor 488 and deoxynucleotidyl transferase (TdT) enzyme to cross-link fluorophore to EdU. Slides then underwent antigen retrieval and regular IHC protocol for immunofluorescent co-labeling. For quantification, 4x images were obtained from caudal and rostral section of CA1 region separated by 800um and averaged together. Images were analyzed using Qupath for TUNEL+ cell counts.

### FerroOrange Assay

After euthanasia, hippocampus was dissected out and 2 right hippocampi were combined for HI samples, 2 left hippocampi for contralateral samples and both right/left hippocampi combined for sham animals. Tissue was dissociated using Miltenyi Adult Brain dissociation kit and Octodissociator. Cell suspension was then filtered through 70um filter and spun down at 300g for 10 minutes to pellet cells. Cells then incubated in red blood cell lysis solution to remove red blood cells, washed and spun down again at 300g for 10 minutes. Cells then resuspended in PBS/1%BSA at 1×104 cells/uL. 100uL added to warmed NeuroMACs media with NB21 supplement with 1:1000 dilution of FerroOrange dye (Thermofisher Cat. SCT210) with control samples with no dye. Cells incubated in dye at 37C for 30min and then washed in PBS two times and resuspended in filtered FACS buffer (PBS, 1% BSA, 25mM HEPES, DNAse). Cells were analyzed on a WOLF Nanocellect.

### Fluorescence activated cell sorting (FACS) isolation

Using cell dissociation protocol described above. After RBC lysis, cells were filtered through 40um strainer to obtain single cell suspension. After obtaining single cell suspension, cells were incubated in PE-Cy5 anti-CD11b antibody (Biolegend) for 10minutes at 4C in PBS/1%BSA. Cells then washed and stained with Sytox-488 (ThermoFisher S34860) to identify dead cells. Cell suspensions and unstained controls were then resuspended in FACS buffer. CD11b+ cells were sorted using the WOLF Nanocellect flow cytometer. 2-4×10^4^ cells collected per sample. Isolated CD11b+ cells were spun down for RNA isolation.

### Quantitative polymerase chain reaction (QPCR)

RNA isolated from samples using Qiagen RNAeasy kit. RNA converted to cDNA using Applied Biosystem reverse transcriptase kit (Cat. 4368814). Subsequently, target genes pre-amplified using Taqman preamplification kit. RT-PCR then run using Taqman assays (see Supplementary Table 1) and run on Applied Biosystem Quant 7 QPCR system.

### Mass spectrometry

Hippocampal tissue was dissected out, flash frozen and stored at -80C. Tissue was sent to Eicosanoid Core at Vanderbilt University Medical Center on dry ice. Frozen tissue was weighed and immediately added to 0.8mL ice-cold chloroform:methanol (2:1) containing 0.01% BHT. The tissue was homogenized with a hand-held sonicator and the sample tube flushed with stream of nitrogen 30–60 s to remove air and then capped. The tubes are shaken at 22–25°C for 1hr. After shaking, 0.3mL aqueous NaCl (0.9%) was added, and the samples were vortexed for 1 min. Centrifugation for 10 min in a tabletop centrifuge separated the aqueous and organic layers. After centrifugation, the top aqueous layer was discarded. The organic layer was evaporated under a stream of nitrogen until dry. Lipids were resuspended in 0.3mL methanol containing 0.005% BHT. 0.3mL Aqueous KOH (15%, wt/vol) was added to the solution. The tube was vortexed, purged with nitrogen, and capped. Sample tubes were incubated at 37°C for 20 min. After incubation, each sample was adjusted to pH 3 with 1N HCl and diluted to a total volume of 1.2 mL with DI water.

The internal standard [^2^H_4_]-15-F_2t_-IsoP was added before extraction on an Oasis MAX uElution plate (Waters Corp., Milford, MA). Sample wells were first washed with methanol (200uL) followed by 25% methanol in water (200uL). The sample was then loaded into the well and washed with 600uL 25% methanol. Eicosanoids were eluted from the plate with 30uL 2-propanol/acetonitrile (50/50, v/v) containing 5% formic acid into a 96-well elution plate containing 30uL water in each well.

Samples were analyzed on a Waters Xevo TQ-XS triple quadrupole mass spectrometer connected to a Waters Acquity I-Class UPLC (Waters Corp., Milford, MA USA). Separation of analytes is obtained using an Acquity PFP column (2.1 × 100mm) with mobile phase A being 0.01% formic acid in water and mobile phase B acetonitrile. Eicosanoids are separated using a gradient elution beginning with 30% B going to 95% B over 8 minutes at a flow rate of 0.30mL/min.

### Statistical Analysis

All analysis was conducted using Graphpad Prism (version 10). Two-way ANOVA tests with Tukey post-hoc multiple comparison analysis was performed to analyze effect from injury (ipsilateral HI, contralateral hypoxia and sham), time post-HI. Two-way ANOVA analysis for For flow cytometry and mass spectroscopy data, matching was performed for sham and HI mice from same litter to control for possible variability between litters. Error bars in the graphs and values following ± symbol in text represent standard deviations.

## Results

### Timeline of cell death following neonatal HI

Neonatal mice underwent modified Vannucci protocol to induce a right sided ipsilateral hypoxic-ischemic brain injury at post-natal day 9 (PND9), with the contralateral hippocampus being exposed to transient hypoxia only TUNEL staining was used to track cell death, which labels DNA double-strand breaks in cells undergoing death from multiple pathways including apoptosis, ferroptosis, or oxidative stress.

Analysis of temporal changes to TUNEL staining focused on the hippocampus since this region showed most consistent injury between animals as the variability of injury in the cortex and striatum would make interpretation of bulk tissue biochemical assays difficult. Quantitative analysis of TUNEL+ cells showed a 6-fold increase in the ipsilateral HI hippocampus compared to sham control at 6h post-HI. In the HI hippocampus, TUNEL staining peaked at 24h post-HI with normalization by 7 days post-HI, when compared to the contralateral hippocampus and sham control hippocampus (Fig.1A-D,A’-D’,A”-D”). Two-way ANOVA analysis evaluating time and injury showed significant effects of both injury (p<0.0001, ), time (p=0.0001) and interaction between the two variables (p<0.0001). At 6h post-HI,TUNEL+ cell density (cells/mm^2^) in the HI hippocampus was significantly elevated compared to sham control (195.3±104.7 vs 32.7±6.5, p=0.0290). When compared to sham controls, peak difference in TUNEL+ cells in the HI hippocampus occurred at 24h post-HI (477.4±143.4 vs 16.2±7.1, p<0.0001) with continued significant increase at 72h post-HI (412.8±143.4 vs 52.8±35.7, p<0.0001) (Fig.1E,E’). By 7d post-HI no difference in TUNEL staining was noted in HI hippocampus compared to sham. No difference in TUNEL staining was noted in the contralateral hippocampus at any time point. Of note, the majority of TUNEL+ cells were located within the pyramidal cell layers of the HI hippocampus, whereas the dentate gyrus had less TUNEL staining in comparison (Fig.1B”).

**Figure 1.**
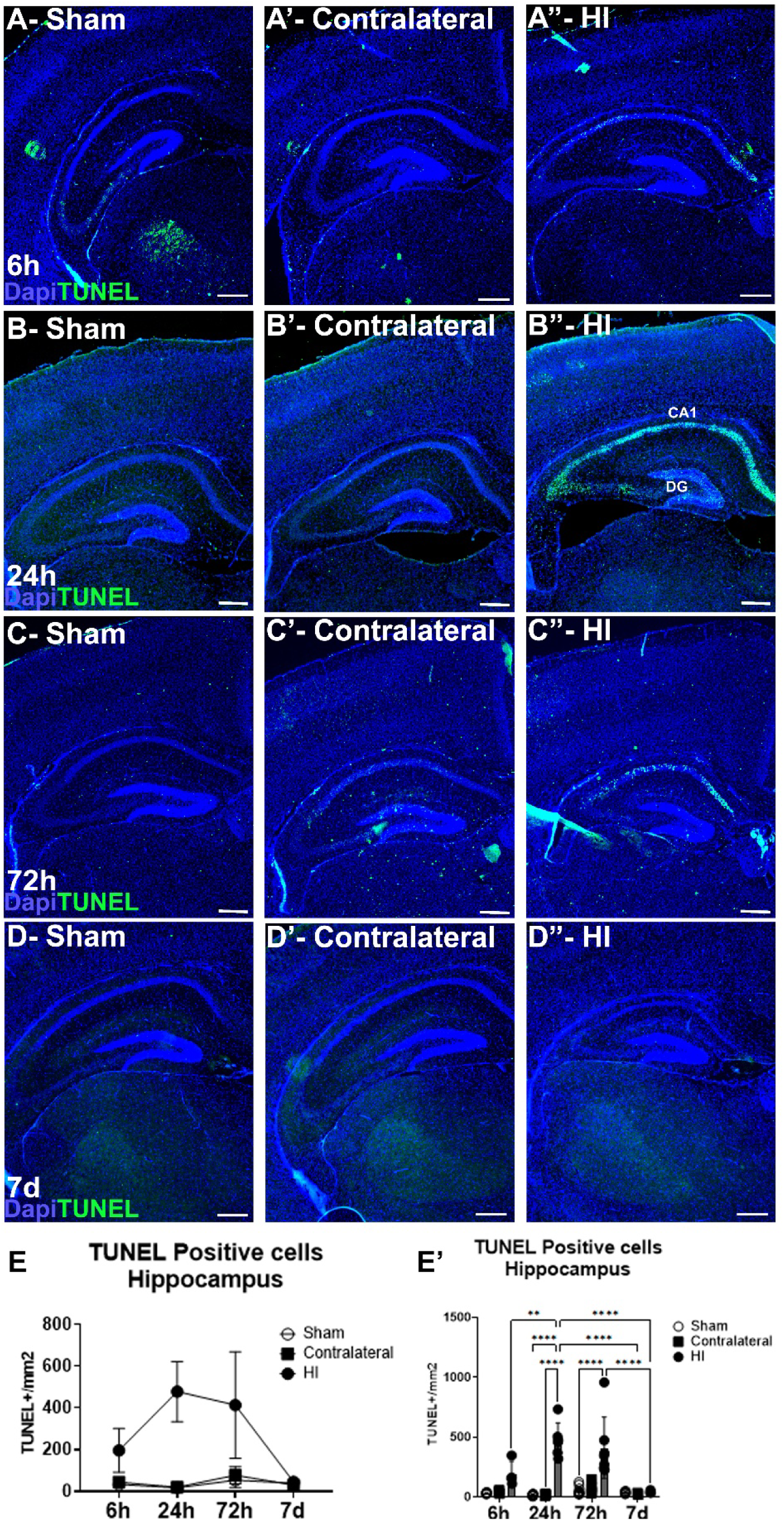
Timeline of TUNEL staining post-HI. TUNEL staining in sham (A-D), contralateral (A’-D’) and HI (A”-D”) hippocampus at 6h (A), 24h (B), 72h (C) and 7d (D) post-HI. (B”) TUNEL staining peaking at 24h post-HI, predominantly in CA1 pyramidal cell layer, with less staining in dentate gyrus (DG). (E,E’) Quantification of TUNEL+ cells in hippocampus post-HI (E) with post-hoc multiple comparison analysis (E’). 2-way ANOVA analysis (injury DF=2, time DF=3, injury*time DF=6) with Tukey post-hoc multiple comparison, **p<0.01, ****p<0.0001 (n=3-6). Scale bar = 100um.

### Elevated labile ferrous iron following neonatal HI

Prior research using the Perls’ stain has shown that neonatal HI brain injury is associated with deposition of iron in areas of ischemic injury.^14,17^ While this stain identifies all non-heme iron, the toxic effects of iron on oxidative stress and cell death are mediated by the free or labile iron pools, with free ferrous iron being most toxic. Thus, the effects of hypoxia and ischemia on intracellular labile iron pools (LIP) has not been extensively studied in a direct manner. To assess changes in ferrous LIP following HI injury, flow cytometry was performed on dissociated hippocampal cells incubated in FerroOrange (FeO) dye, which unlike Perls’ stain, specifically detects labile ferrous iron (Fig.2A,B). Analysis of FeO+ cells in sham, hypoxia alone (contralateral) and HI hippocampus showed an effect from injury (p<0.0001), time (p=0.0017) and interaction between the variables (p=0.016) when analyzed by 2-way ANOVA indicating that cells in the ipsilateral HI hippocampus had elevated LIP (Fig.2C-D). Interestingly, intracellular LIP was maximal at the 6h time-point, with 19.3±3.5% of cells being FeO+ in the HI hippocampus, which was1.74-fold higher compared to cells from the sham hippocampus (11.1±.6, p<0.0001). While at 24h post-HI, the relative number of FeO+ cells in HI hippocampus decreased to 14.5±1.9%, there was still a 2.29-fold increase in FeO+ cells compared to sham control (6.3±1.4, p<0.0001). Finally, 10.1±2.4% of cells were FeO+ in the HI hippocampus at 72h post-HI which was 1.64-fold higher compared to sham control (6.2±2.6, p=0.001). Interestingly, there was a 57% decrease in the percentage of FeO+ cells over time in the sham controls between PND9 and PND12 (p=0.01) (Fig.2C-D). There was no significant different in FeO+ cells in the contralateral hippocampus when compared to sham.

**Figure 2.**
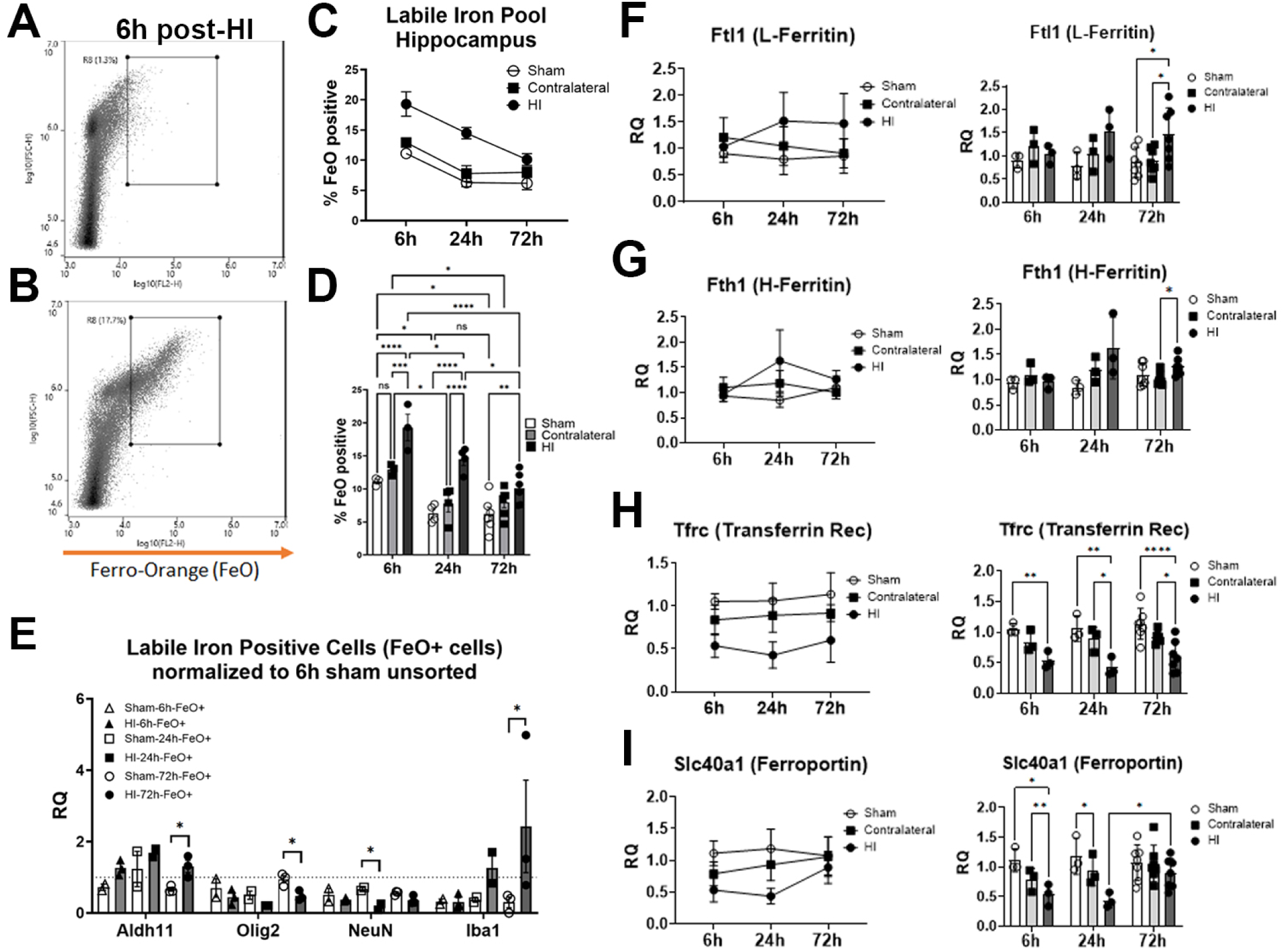
Elevated post-HI intracellular labile iron in hippocampal cells is accompanied by dysregulated expression of iron metabolism genes. Flow cytometry showing FerroOrange signal from dissociated hippocampal unstained control (A) and FerroOrange stained cells (B) at 6h post-HI. (C) Quantification of FerroOrange positive cells from sham, contralateral and HI dissociated hippocampal cells. (D) Bar graph of summarized plot showing Tukey post-hoc multiple comparison analysis (n=3-5). (E) Relative quantification of cell specific gene expression for astrocytes (Aldh1l1), oligodendrocytes (Olig2), neurons (NeuN) and microglia/macrophages (Iba1) from FerroOrange sorted cells (n=2-3). (F-I) Relative quantification of gene expression in bulk hippocampal cells of L-ferritin (F), H-ferritin (G), transferrin receptor (H) and ferroportin (I) with bar graphs showing post-hoc multiple comparison analysis. Samples were normalized to sham 6h sample (n=3-6). Two-way ANOVA analysis (injury DF=2, time DF=2, injury*time DF=4, matched by litter mates) with Tukey post-hoc multiple comparison, *p<0.05, ***p<0.001, ****p<0.0001.

Next, sorted FeO+ cells from the sham or HI (ipsilateral) hippocampus were analyzed by qPCR to identify which cell types predominated (Fig. 2E). This showed significant enrichment of astrocyte-specific genes (Aldh1l1) early after HI and microglia/macrophages markers (Iba1) which was significant at 72h post-HI. Markers of neurons (NeuN) and oligodendrocytes (Olig2) gene expression were relatively depleted in FeO+ cells, but only significantly so at 24h and 72h post-HI, respectively. Data was analyzed by 2-way ANOVA with Aldh1l1, Olig2 and NeunN showing effects from injury (Aldh1l1: p=0.008, Olig2: p=p=0.004, NeuN: p=0.011) but no effect from time or interaction between variables. Iba1 did not show an effect from HI, time or interaction, but post-hoc analysis showed a significant effect between HI and sham at 72h post-HI. These findings indicate that intracellular labile iron accumulated within viable astrocytes and microglia. Sorting FeO+ cells from the contralateral hippocampus was not feasible due to technical limitations of sorting time and stability of the FeO signal, so HI and sham samples were prioritized.

### Transcriptional changes to iron regulatory genes indicate neuronal intracellular iron sequestration following neonatal HI

While FerroOrange has been used for *in vitro* studies, its use in dissociated tissue has not previously been tested. Furthermore, sorting FeO+ cells posed technical challenges that limited the number of samples with enough RNA yield to analyze gene expression. Thus, confirming intracellular iron accumulation with corresponding gene expression changes was needed to validate FerroOrange findings. Increases in intracellular iron typically result in decreases to iron uptake genes such as transferrin receptor, which is one of the most sensitive markers of intracellular labile iron changes. In addition, iron storage genes such as H-ferritin and L-ferritin also increase to promote iron storage. Finally, the iron export protein ferroportin (Slc40a1) typically increases when intracellular iron is elevated, but in the setting of iron sequestration, expression of Slc40a1 decreases. In order to correlate FeO changes with gene expression changes, bulk qPCR analysis of dissociated hippocampal cells also showed gene expression changes consistent with increased labile iron and iron sequestration. This included a decrease in Tfrc (transferrin receptor), increased Ftl1 (L-ferritin), transiently increased Fth1 (H-ferritin) and decreased Slc40a1 (ferroportin) gene expression (Fig.2F-I). These changes were evident as early as 6h post-HI and persisted to 72h post-HI. 2-way ANOVA analyses of these genes showed significant effects from injury in Tfrc (p=0.0002), Ftl1 (p=0.026), Fth1 (p=0.04) and Slc40a1 (p=0.035) (Fig.2F-I). Most prominent effects were in HI hippocampus in post-hoc comparison analysis particularly in Ftl1, Tfrc and Slc40a1, but contralateral hypoxia exposed hippocampus also showed changes that were indicative of minor activation of an iron sequestration response (Fig.2F-I). None of the genes showed a significant time effect or interaction between time and injury based on 2-way ANOVA analysis.

Further analysis to see which cells showed gene expression changes was conducted using fluorescent *in situ* labelling with RNAscope at all time-points (Fig.3A-D). This showed elevated Ftl1 gene expression and decreased Tfrc expression within CA1 region post-HI that was significant for HI when analyzed by 2-way ANOVA (Ftl1: p=0.001; Tfrc: p=0.006) also suggesting that these cells had higher labile iron levels (Fig.3E-F). Post-hoc analysis showed that Ftl1 expression was highest in HI hippocampus at 24h post-HI compared to sham. Both Ftl1 and Tfrc staining analysis did not meet significance for time effect (Ftl1: p=0.074; Tfrc: p=0.058), but significant interaction between time and injury (Ftl1: p=0.003; Tfrc: p=0.037). Interestingly, cells with pyknotic nuclei (which are identified by condensed chromatin resulting in small homogenous dapi stain) within the pyramidal layer had very high expression levels of Ftl1 compared to surrounding cells (Fig.3A-D, arrowheads) suggesting that these cells were attempting to respond to the toxic effects of labile iron. Furthermore, cells in the CA1 region of the contralateral hippocampus (hypoxia exposure) also showed elevated Ftl1 staining and Tfrc staining at 24h post-HI.

**Figure 3.**
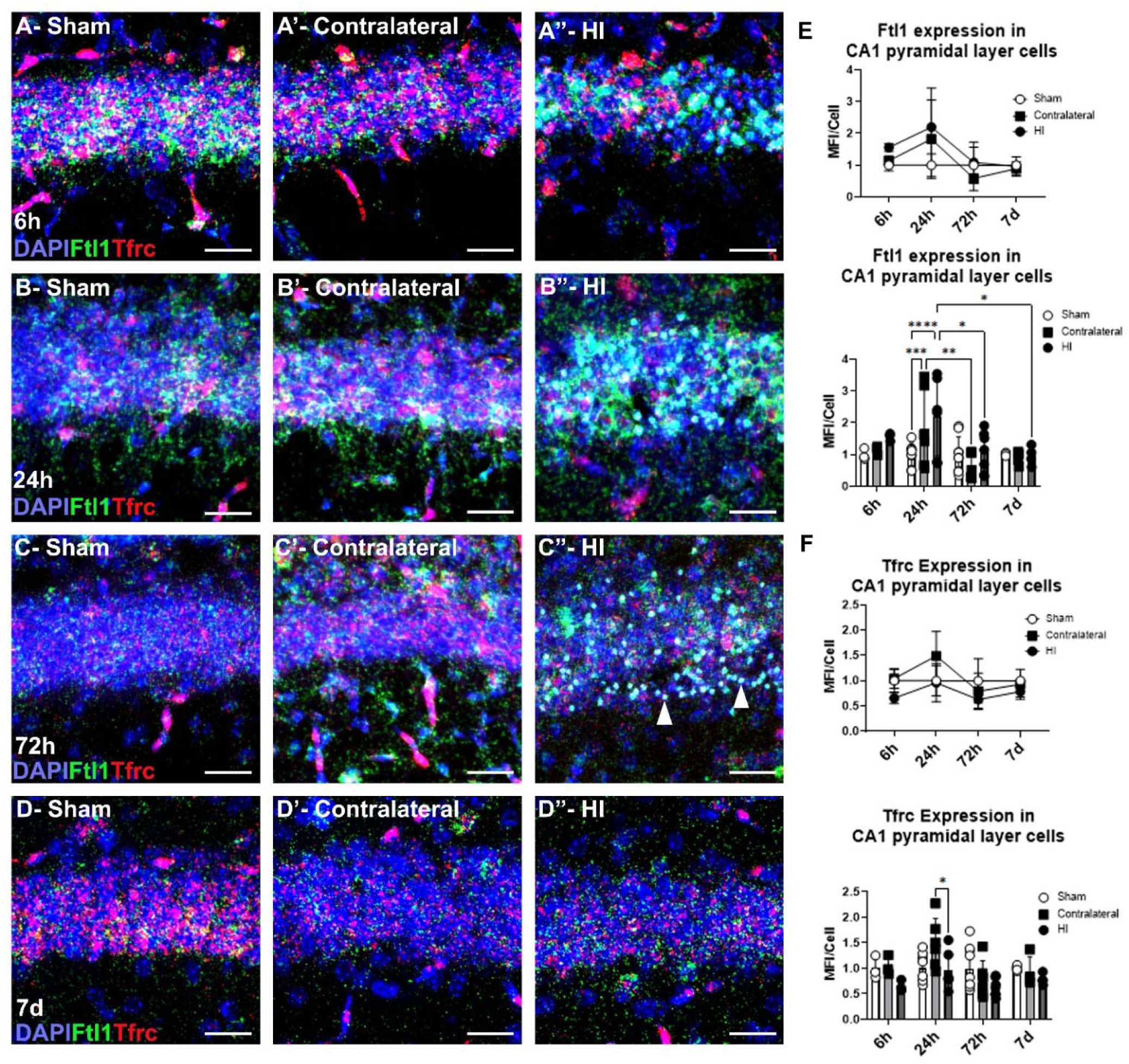
Time-dependent effects on L-ferritin and transferrin receptor gene expression in hippocampal CA1 pyramidal cell layer. (A-D) Fluorescent *in situ* of sham, contralateral and HI at 6h (A), 24h (B), 72h (C) and 7d (D) post-HI showing transcriptional changes to Ftl1 (L-ferritin) and Tfrc (transferrin receptor) as surrogate markers for intracellular iron changes. (C”) Arrowheads identifying pyknotic nuclei that are Ftl1 positive. (E,F) Quantification of mean fluorescent intensity of Ftl1 (E) and Tfrc (F) staining per cell (identified by DAPI stain) normalized to sham controls at each specific time-point, graphs show both summary analysis and bar graph with multiple comparison. Analyzed by 2-way ANOVA (injury DF=2, time DF=3, injury*time DF=6) with Tukey post-hoc multiple comparison, *p<0.05, ***p<0.001, ****p<0.0001 (n=3-6). Scale bar = 20um.

### Increased iron sequestration response primarily in microglia/macrophages

Given the upregulation in Ftl1 gene expression and decrease in labile iron over a 72 hour period, double labeling with L-ferritin and cell specific protein markers was performed to determine which cell types sequester iron post-HI. Increases in L-ferritin protein levels would increase iron storage capacity and decrease labile iron pool levels. Since HI labile iron levels were close to sham levels at 72h, this time-point was initially chosen to look for L-ferritin protein expression. Immunofluorescence at 72h post-HI showed increased L-ferritin positive cells in areas where ischemic injury occurred based on TUNEL staining (Fig.4A,A’). Co-labeling L-ferritin with NeuN (neurons), Gfap (astrocytes) showed minimal colocalization, but there was prominent overlap with Iba1+ cells (microglia/macrophages) (Fig.4B-D). Interestingly, Iba1+/FtL+ cells were located throughout the hippocampus, but their presence in the pyramidal cell layer was of particular interest as they appeared in areas where NeuN staining was lost (Fig.4E,E’).

**Figure 4.**
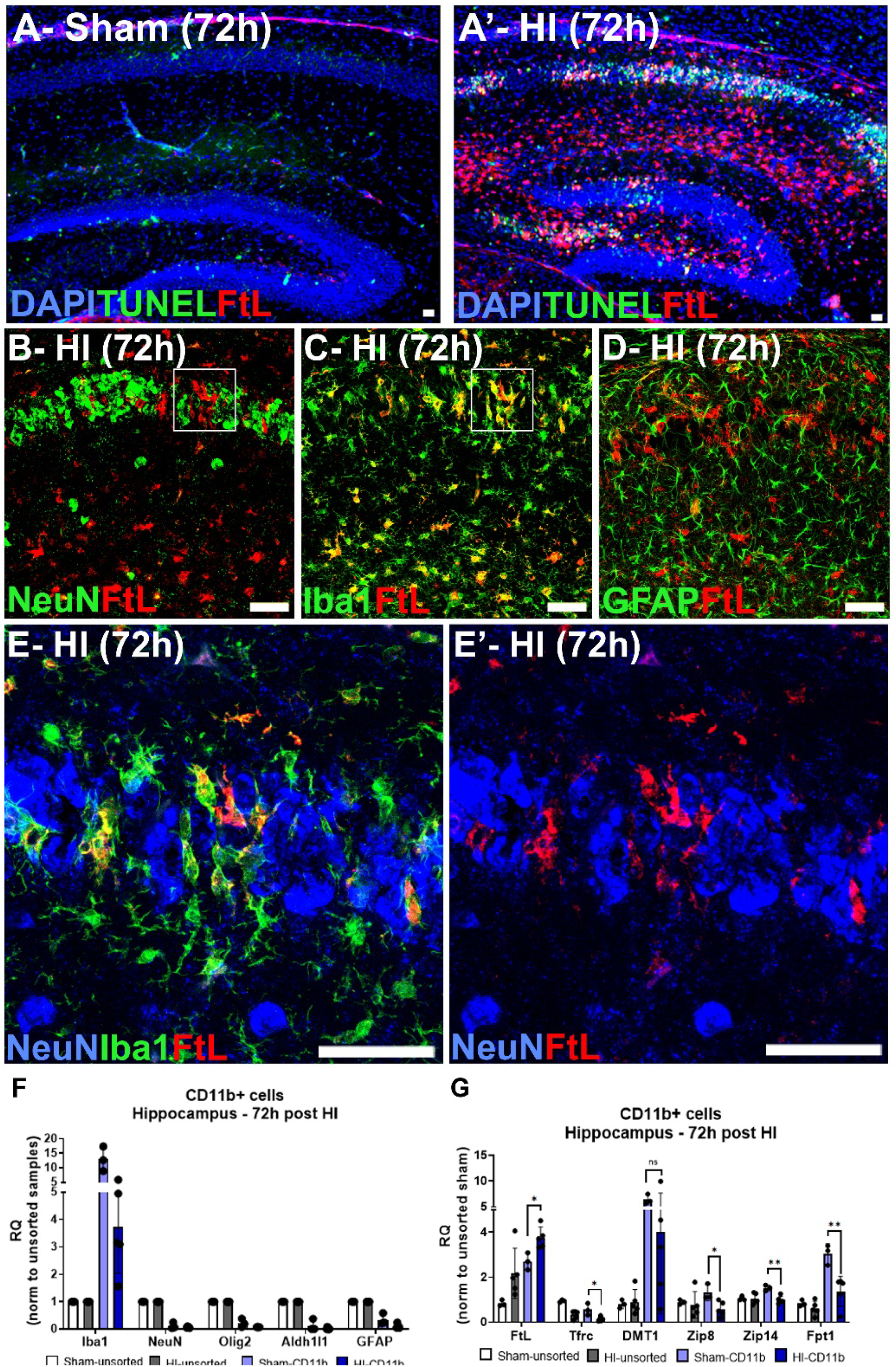
L-ferritin protein co-localizes with microglia/macrophages. (A,A’) L-ferritin (FtL) immunofluorescent staining shows elevated FtL in HI hippocampus in areas of TUNEL staining when compared to sham. (B-D) Colocalization of FtL with NeuN+ neurons (B), Iba1+ microglia/macrophages (C) and GFAP+ astrocytes (D) in hippocampus 72h post-HI showing significant co-localization with Iba1+ cells. (E,E’) Higher magnification image of CA1 pyramidal cell layer in box from B and C showing NeuN, Iba1 and FtL triple-labeling. (F,G) QPCR from sorted CD11b+ cells from sham and HI hippocampus at 72h post-HI showing enrichment of Iba1+ cells in sorted population (F) and gene expression changes of cellular iron regulatory genes. Analysis via t-test comparing sham vs. HI CD11b+ cells, *p<0.05, **p<0.01 (n=3-5). Scale Bar = 20um

In order to validate whether microglia/macrophages were specifically sequestering iron, CD11b+ cells were isolated at 72h post-HI from hippocampus of HI and sham mice via flow cytometry to evaluate changes in expression of genes that regulate iron metabolism. The sorted CD11b+ cells were enriched for microglia/macrophage as expected, since expression of neuronal, astrocytic and oligodendrocyte markers was low, while microglia/macrophage markers were elevated when compared to unsorted samples (Fig.4F). QPCR analysis of CD11b+ sorted cells showed decrease in Tfrc (75% reduction, p=0.014) and Slc40a1 (55% reduction, p=0.009), but increased Ftl1 (40% increase, p=0.018) consistent with increased intracellular iron content. Other ferrous iron import proteins were also evaluated which included Dmt1 (Slc11a2), Zip8 (Slc39a8) and Zip14 (Slc39a14). Gene expression of iron import genes was significantly decreased following HI, except for Dmt1 (Zip8: 55% reduction, p=0.027; Zip14: 35% reduction, p=0.003). Furthermore, CD11b+ cells had relative enrichment of DMT1 gene expression compared to Zip8 and Zip14 gene expression (Fig.4G).

### Microglial/macrophage iron sequestration response peaks at 72h post-HI

Since L-ferritin protein was upregulated in microglia/macrophages, the temporal pattern of microglia/macrophage infiltration and their cellular iron sequestration response was assessed to determine how this response related to labile iron accumulation and cell death. Double label immunofluorescence for FtL1 and Iba1 was performed at 6h, 24h, 72h and 7d post-HI (Fig.5A-D, A’-D’, A”-D”). All imaging and analysis were completed by examining the CA1 region as this was the primary region of cell death occurrence. Between P9 and P12-P16, there was an increase in Iba1+ cells from 4% to 12% of total cells consistent with previously observed microglial expansion during the first 2 post-natal weeks. Iba1+ cells in CA1 region of HI hippocampus increased up to 26% of total cells (p<0.0001 when compared to sham) with statistical significance from injury (p<0.0001), time (p<0.0001) and the interaction of these variables (p<0.0001) when analyzed by 2-way ANOVA (Fig.5E,E’). In sections from sham controls, 18-24% of Iba1+ cells were FtL+ at all time-points. However, in the HI hippocampal CA1 region, 26.8% of Iba1+ cells were FtL+ at 6h, peaking to 82.7% at 72h, and decreasing to 37.5% of total Iba1+ cells by 7d post-HI (Fig.5F,F’). Interestingly, the contralateral hippocampus showed a transient increase in L-ferritin immunoreactivity in Iba1+ cells 24h post-HI (p=0.046) that is apparent on imaging along with morphological changes to microglia (increased cell soma with thicker processes), with subsequent normalization by 72h post-HI. When analyzed by 2-way ANOVA, Iba1+/FtL+ double positive cells showed a significant effect of injury (p<0.0001), time (p=0.0001) and interaction between the variables (p=0.0003) (Fig.5A-F). These findings confirm that microglia/macrophages activate an iron sequestration response that is activated by hypoxia, but persists and is amplified by the presence of neuronal cell death.

**Figure 5.**
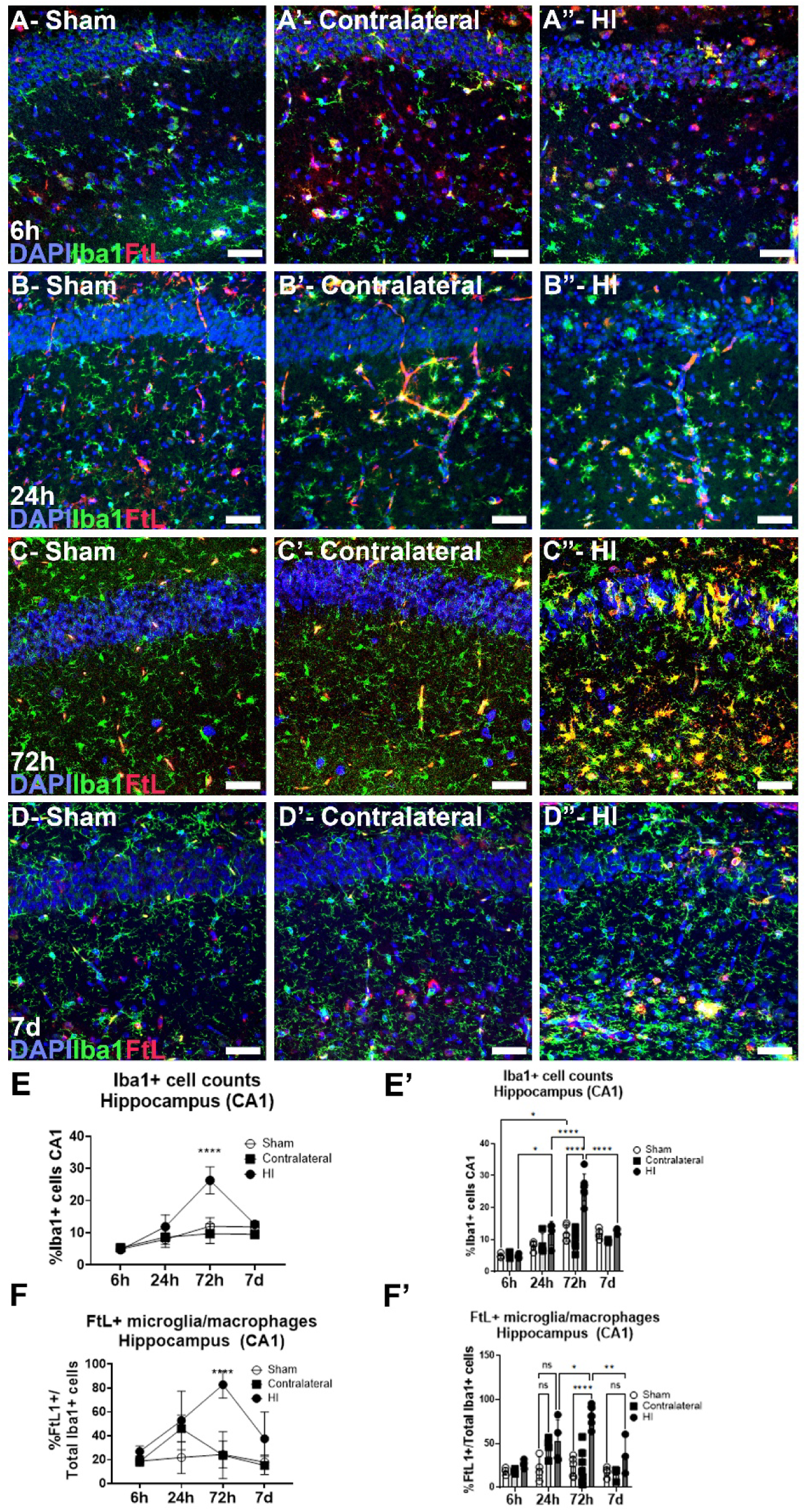
L-ferritin positive microglia/macrophages show peak response at 72h post-HI. Immunofluorescent staining of L-ferritin (FtL) and Iba1 (microglia/macrophages) in sham (A-D), contralateral (A’-D’) and HI (A”-D”) hippocampus at 6h (A), 24h (B), 72h (C) and 7d (D) post-HI. Scale bar = 20um. Quantification of percent of Iba1+ cells in CA1 region of hippocampus with bar graph showing post-hoc multiple comparison analysis (E,E’). Quantification of percent of Iba1+ cells that are FtL+ over time post-HI with bar graph showing post-hoc multiple comparison analysis (F,F’). Analyzed via 2-way ANOVA (injury DF=2, time DF=3, injury*time DF=6) with Tukey post-hoc analysis (n=3-7), *p<0.05, **p<0.01, ****p<0.0001

### Early lipid peroxidation changes post-HI

Iron mediated production of hydroxyl radicals can oxidize and damage tissue. Specifically, polyunsaturated fatty acids (PUFAs) are susceptible to oxidation, which is a key feature of ferroptosis.^8^ Non-enzymatically oxidized PUFAs can form metabolites called isoprostanes (derived from arachidonic acid) and neuroprostanes (derived from docosahexaenoic acid) (Fig.6A).^21,22^ There are multiple different neuroprostane and isoprostane species but the most prominent ones in the brain include the 4-series neuroprostanes (4-F_4t_-NP) and the 15-series isoprostane (15-F_2t_-IsoP, often referred to as 8-iso-PGF_2α_).^23,24^ Isoprostane and neuroprostane species were measured in hippocampus at 6h, 24h and 72h post-HI. HI hippocampus showed a 1.95 fold increase in 4-F4t-NP levels at 24h (p=0.043) that declined by 72h post-HI when compared to sham hippocampus. Interestingly, the contralateral hippocampus also showed a trend towards elevated 4-F_4t_-NP levels at 24h as well (1.8 fold increase, p=0.087) (Fig.6B-C). Overall, when analyzed by 2-way ANOVA, there was significant effect from time (p=0.0141) and a significant interaction between injury and time (p=0.0324), but there were no significant effects due to injury which is likely because there was only a transient effect at 24h post-HI. In contrast, 15-F_2t_-IsoP levels did not exhibit any significant effect of injury, time or interaction between the two variables. The oxidation of DHA was validated by immunofluorescent staining for CEP (carboxyethylpyrrole), which is a split fragment product of DHA non-enzymatic oxidation. CEP staining was noted in TUNEL positive cells within the CA1 region of the hippocampus at 24h post-HI (Fig.6D). This indicates that non-enzymatic oxidation of docosahexaenoic acid (DHA) occurs within the initial 24h post-HI but normalizes by 72h, while arachidonic acid (ARA) oxidation does not significantly change during the acute injury period when neuronal cell death occurs. Furthermore, markers of oxidized DHA were evident in TUNEL positive cells indicating their association with cell injury.

**Figure 6.**
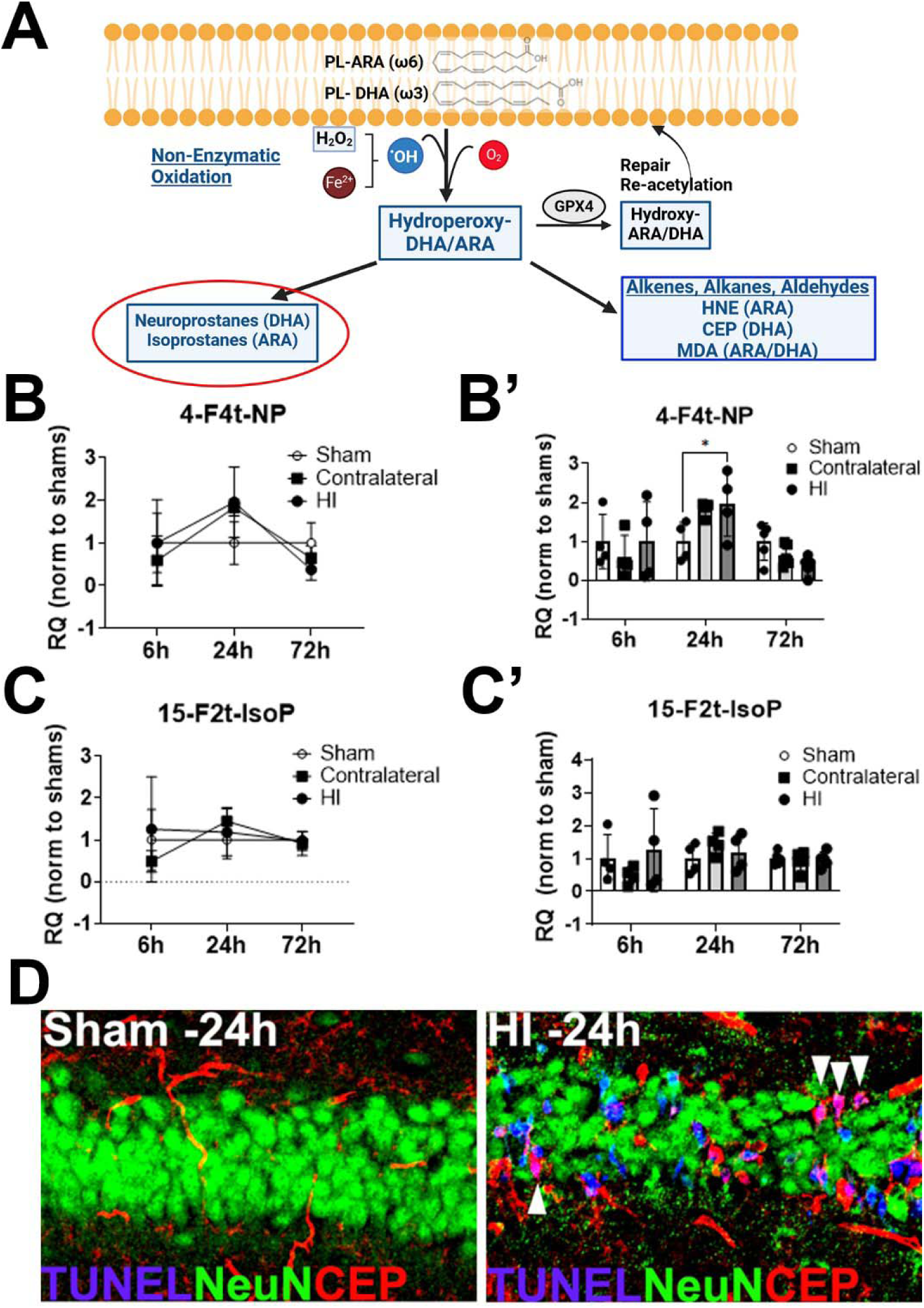
Markers of lipid peroxidation show elevated DHA oxidation 24h post-HI. (A) Schematic showing by-products of non-enzymatic oxidation of the primary PUFAs in the brain (ARA and DHA), which results in neuroprostanes (DHA derived) and isoprostanes (ARA derived). (B,B’) Mass spectroscopy quantification of 4-series neuroprostanes (4-F4t-NP) at 6h, 24h and 72h post-HI with bar graph showing post-hoc multiple comparison analysis. (C,C’) Mass spectroscopy quantification of primary 8-series isoprostane, 15-F2t-IsoP, at 6h, 24h and 72h post-HI with bar graph showing post-hoc multiple comparison analysis. Two-way ANOVA analysis (injury DF=2, time DF=2, injury*time DF=4, matched by litter mates) and Tukey post-hoc multiple comparison, *p<0.05 (n=4-5). (D) Immunofluorescent staining of TUNEL, NeuN (neurons) and CEP (oxidized DHA) staining 24h post-HI.

### Ferroptosis gene expression markers change following neonatal HI

Since lipid peroxidation is the downstream effector of cell death by ferroptosis, the expression of select ferroptosis genes over the first 72 hours post-injury were assessed using bulk qPCR of dissociated hippocampal cells (Fig.7A). ^25–27^ The evaluated genes included Ncoa4, which drives the increase in LIP from ferritin degradation, Acsl4, which is involved in the production of polyunsaturated fatty acid lipids, and Gpx4 and Slc7a11, which are responsible for lipid peroxide neutralization (Fig.7B-E). Ncoa4 showed a decreased in expression with significant effect from injury (p=0.048) with post-hoc analysis showing significant effect at 24h (p=0.033), but no effect from time (p=0.053) or interaction (p=0.27). Analysis of Acsl4 showed a significant effect from injury (p=0.014) causing a decrease in Acsl4. Post-hoc analysis showed significant decreases of Acsl4 in HI tissue compared to contralateral tissue at 24h (p=0.016) and 72h post-HI (p=0.006). Gpx4 gene expression showed a significant effect of time (p=0.034), but no effect from injury or interaction. Finally, analysis of Slc7a11 showed a significant effect from time (p=0.015) and interaction effect (p=0.029), no effect from injury. However, HI tissue showed significant increase compared to sham at 72h post-HI (p=0.009) on post-hoc analysis. These findings show that viable cells from the HI hippocampus are potentially inducing gene expression changes that are anti-ferroptotic as decreases in Acsl4 and Ncoa4 while increases in Slc7a11 would provide protective effects.

**Figure 7.**
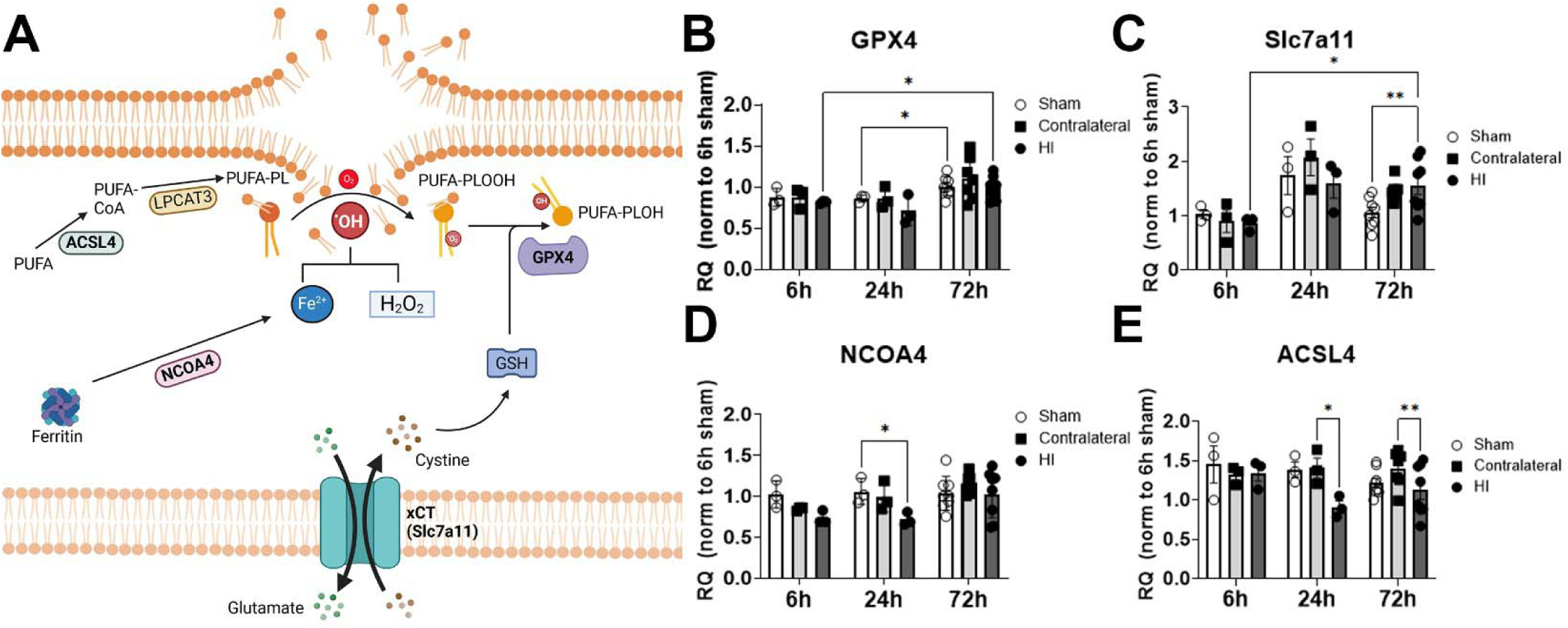
Gene expression changes in markers of ferroptosis post-HI. (A) Schematic showing proteins involved in ferroptosis, namely ACSL4, NCOA4, GPX4 and Slc7a11. (B-E) Relative quantification of gene expression of NCOA4 (B), ACSL4 (C), GPX4 (D) and Slc7a11 (E) at 6h, 24h and 72h post-HI. All samples normalized to a 6h sham control sample. Two-way ANOVA (injury DF=2, time DF=2, injury*time DF=4, matched by litter mates) with Tukey post-hoc multiple comparison analysis (n=3-6).

### Increased neuronal Gpx4 expression following HI

While there was no change in gene expression of Gpx4, we have shown elevated neuroprostanes that normalize by 72h. The decline in 4-F4t-NP between 24h and 72h post-HI would indicate that there is a mechanism either due to decreased availability of PUFAs or increased clearance of lipid peroxides. Lipid peroxides are neutralized by Gpx4, so while there was no change in the transcript level in bulk hippocampal cells, we evaluated if there was any cell specific differences at the protein level. Since DHA is enriched within neurons, we used immunofluorescence to assess Gpx4 expression within NeuN+ neurons (Fig8A-D,A’-D’). NeuN+ CA1 pyramidal neurons showed elevated Gpx4 expression that was statistically significant at 72h post-HI when compared to sham (114±17.7 vs 75.6±6.7, p=0.035), which indicates that remaining neurons following HI might be protected by elevated Gpx4. Overall, when examined by 2-way ANOVA, Gpx4 mean fluorescent intensity in NeuN+ cells showed a statistically significant effect of injury (p=.042) and time (p=0.004), but no interaction between the two variables (Fig.8E,E’). Although Gpx4 protein expression was most prominent in neuronal cells, as evidenced by co-labeling with NeuN+ cells, colocalization was also observed in Iba1+ cells particularly those that infiltrated into the pyramidal layer following HI (Fig.8F-H). The contralateral hippocampus showed no difference in Gpx4 immunoreactivity within neurons.

**Figure 8.**
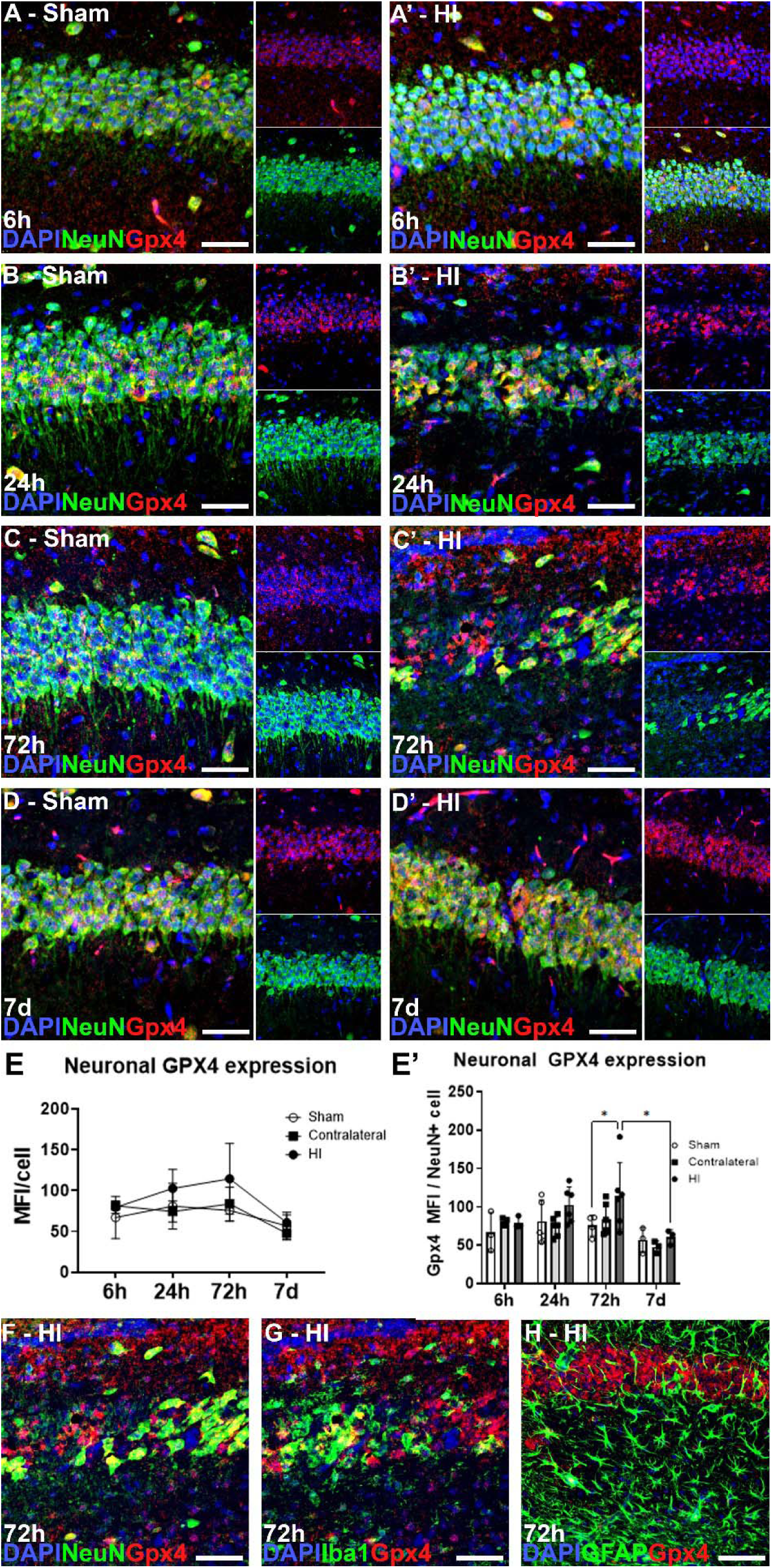
Neuronal upregulation of GPX4 peaks at 72h post-HI. (A-D) Immunofluorescent co-labeling of NeuN and Gpx4 at 6h, 24h, 72h and 7d post-HI. (E,E’) Quantification of mean fluorescent intensity of Gpx4 staining in NeuN positive cells. Bar graph shows post-hoc multiple comparison analysis. (F-H) Co-localization of Gpx4 with NeuN+ neurons, Iba1+ microglia/macrophages and GFAP+ astrocytes which shows predominant colocalization with NeuN+ cells. Two-way ANOVA analysis (injury DF=2, time DF=3, injury*time DF=6) with Tukey post-hoc multiple comparison, *p<0.05 (n=3-6). Scale bar = 20um.

## Discussion

Herein, we identify the temporal changes to intracellular iron regulation and lipid peroxidation and correlate it with cell death in the hippocampus following neonatal hypoxic-ischemic injury to determine the relationship between cell specific responses iron dysregulation and lipid peroxidation and neuronal cell death.

We showed that the labile iron level in hippocampal cells was maximally increased at 6h post-HI with steady decline at 24h and 72h post-HI. Labile iron levels remained elevated in hippocampal cells by 72h post-HI although by only marginal amount. The elevated labile iron levels precede the increase in TUNEL staining within the hippocampus and the decline in TUNEL staining after 24h correlates with a decrease in labile iron levels.

Sorting of FeO+ cells revealed increases mostly in microglia/macrophages and to lesser extent in astrocyte markers gene expression by 72h, while markers of oligodendrocytes and neurons appeared to be downregulated. However, given sorted cells may miss dying neurons or injured cells, *in situ* labeling was performed to evaluate cell specific changes to iron regulation in areas of injury. Indeed, we did note cells in the pyramidal layer upregulating Ftl expression and downregulating Tfrc with many condensed nuclei (pyknotic cells) showing markedly elevated Ftl expression. Gene expression changes in unsorted hippocampal cells were also consistent with cellular iron accumulation as evidenced by increased in Ftl1 (L-ferritin), decreased Tfrc (transferrin receptor) and decreased Slc40a1 (ferroportin). We further showed that cells with the most robust iron sequestration response at 72h post-HI appeared to be microglia/macrophages as they showed the most prominent increase in L-ferritin protein levels.

Next, we showed that lipid peroxidation is a more acute response with specifically DHA showing more susceptibility to oxidation compared to ARA following HI. Prior work in humans has also shown that isoprostanes and neuroprostanes can be elevated following HI, however, our data show the time-line of these changes as well as their relationship to TUNEL reactivity and labile iron changes.^28,29^ Interestingly, the peak oxidation of DHA aligns with peak changes in TUNEL staining and is preceded by maximal labile iron levels, which is indicative that acute (<24h) elevations in labile iron may be key contributor to lipid peroxidation. Oxidation of DHA is an important pathologic mechanism in neonatal HI as published literature has shown that supplementation of DHA prior to injury has therapeutic benefit.^30^

Since labile iron levels and neuroprostane levels decline by 72h, we also show potential mechanisms by which this may occur. Infiltration of microglia/macrophages peaks around 72h post-HI and this is associated with upregulation of L-ferritin protein expression along with gene expression changes that indicate a cellular iron sequestration response characterized by increased Flt1, decreased Tfrc and decreased Slc40a1. The correlation between this response and subsequent decline in lipid peroxidation, TUNEL+ density and labile iron levels suggests that this may be a mechanism to decrease toxic effects of labile iron by sequestration in ferritin. Indeed, recent studies have shown that microglia depletion prior to HI can result in worse injury, although this effect appears to be sex specific.^31,32^ The temporal infiltration of microglia/macrophages into areas of ischemic injury is similar to prior studies.^33^ We did not observe any sex specific changes in molecular markers but future studies that manipulate these molecular mechanisms would need to be powered to identify any sex specific effects.

Finally, we also observed temporal changes in Gpx4 protein levels following HI with an increased in Gpx4 immunoreactivity in NeuN+ cells in the CA1 region following HI that peaked at 24-72h post-HI and normalized by 72h post-HI. The increase in Gpx4 protein expression also correlates with a decline in isoprostane and neuroprostane levels between 24h and 72h post-HI. This suggests that there is a compensatory increase in Gpx4 immunoreactivity following HI that may be neuroprotective. Indeed, cells that maintained NeuN+ staining showed elevated Gpx4 expression compared to NeuN+ cells in the sham controls. Interestingly, bulk QPCR analysis showed no changes in Gpx4 gene expression, but this may result from loss of neurons at this timepoint. Prior work looking at ferroptosis and Gpx4 staining showed decreases in protein and gene expression at 24h and 72h post-HI, however these studies were performed using bulk western blots or bulk QPCR analysis.^6,34–38^ This prevents the possibility of looking for spatial and cell specific changes. Differences in injury severity could also explain differences in markers of ferroptosis observed in this study compared to other studies. Extensive neuronal cell death from severe injury could account for the decline in Gpx4 protein levels when compared to this study which had a milder injury and analyzed dissociated cells which likely biases analysis towards viable cells that are able to tolerate tissue dissociation.

Importantly, changes in the contralateral hippocampus (hypoxia exposed) revealed interesting changes that showed potential gene expression changes to augment iron sequestration within hippocampal cells with transient increase in L-ferritin in microglia. However, there was no increase in labile iron pools based on FeO analysis. This is potentially a response from hypoxia as activation of Hif1a and Hif2a are known to target iron regulatory genes like Tfrc and ferritin.^39,40^ Interestingly, the contralateral hippocampus also showed a trend towards elevated neuroprostanes compared to sham as well. It is not clear if the lipid peroxidation in the contralateral hippocampus is derived from the presence of ischemic tissue in the HI hippocampus or a direct effect from hypoxia. If just a result of hypoxia, it is possible that lipid peroxidation is not sufficient to induce cell death but requires additional stressors to cause cell death. Oxidation of lipids has been seen in hypoxia, but isoprostanes and neuroprostanes have not been measured in this context.^41,42^ Lastly, this analysis identified lipid peroxidation in whole tissue, but there may be cell specific differences in DHA peroxidation that lead to neuronal injury. Thus, iron metabolism and lipid peroxidation may also be affected by hypoxia alone, but the presence of neuronal cell death appears to specifically drive labile iron increases that may produce more toxic effects of lipid oxidation and oxidative stress that potentially overwhelm endogenous protective mechanisms and contribute to ongoing cell injury. Interestingly, transient hypoxic pre-conditioning is known to provide a protective response against ischemic injury and oxidative stress.^43–46^ Thus, priming cells for iron sequestration may be one mechanism by which pre-conditioning has therapeutic benefit.

Overall, this data shows the temporal changes in labile iron, lipid peroxidation and peroxidase defense systems. These results show that labile iron and lipid peroxidation increase very early following HI which confirm their toxic effects as acute pathologic responses but there appears to be a compensatory response to augment iron storage through upregulation of ferritin, particularly in phagocytic cells, and neutralization of lipid peroxyl radicals through neuronal Gpx4 upregulation (Fig.9). The extent to which this indicates that ferroptosis is occurring is difficult to quantify. However, oxidative stress and lipid peroxidation can contribute to other forms of cell death that include apoptosis. Thus, the toxic effects of labile iron and lipid peroxidation are likely contributing to cell death through multiple mechanisms.

**Figure 9.**
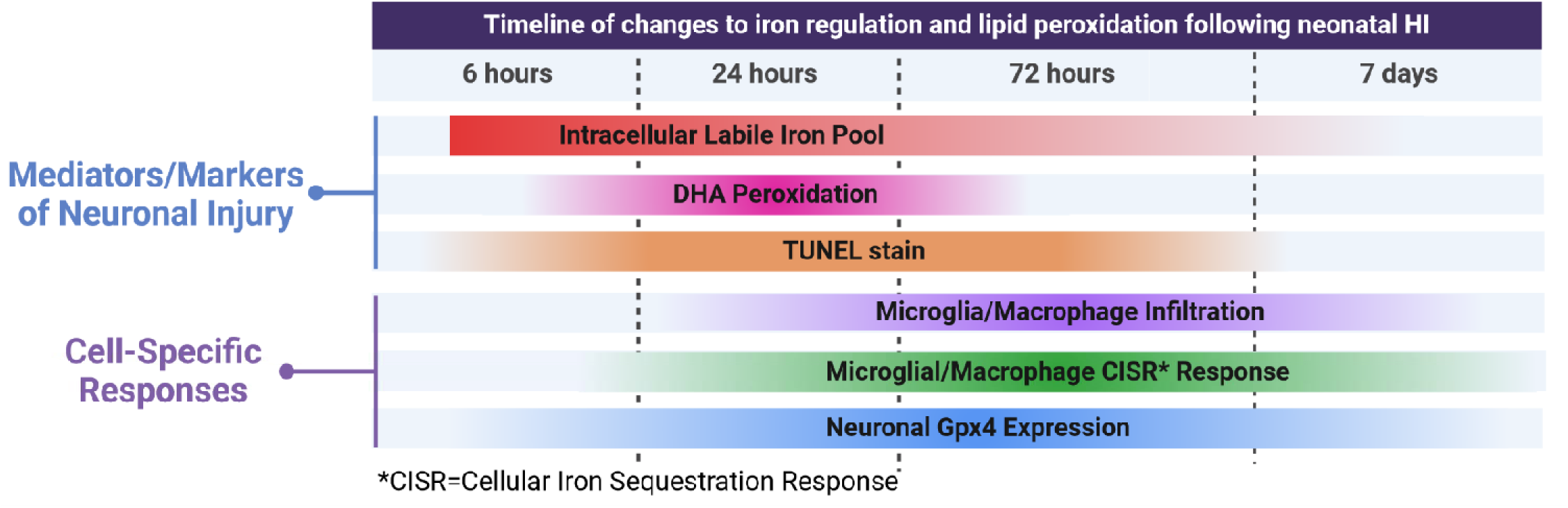
Temporal changes for individual molecular markers of iron regulation and lipid peroxidation and the cell-specific responses in HI hippocampus. Summary of findings from this study showing timeline of molecular changes post-HI. Early increase in intracellular labile iron is followed by peak in lipid peroxidation, which coincides with peak levels of TUNEL staining. These markers of cell injury start to decline as L-ferritin positive microglia/macrophages infiltrate HI tissue and NeuN+ neurons upregulate Gpx4 protein levels.

Some of the caveats to our study include small sample sizes with our analysis of FeO+ sorted cells, but this is because the FerroOrange dye signal is not particularly stable for a prolonged period which affects total sample yield during sorting. However, we tried to address this by using our *in situ* analysis, bulk QPCR analysis and immunofluorescence to show that these indirect measures of cellular iron status correlated with FeO flow cytometry data. In addition, although not powered to evaluate for sex specific effects, we did not observe any sex related differences in these markers. However, previous studies have shown sex related differences in ferroptosis susceptibility and iron toxicity which suggests that any interventions targeting lipid peroxidation or iron toxicity may have sex specific effects.^47,48^ The data showing a cellular iron sequestration response in Iba1+ and isolated CD11b+ cells does not distinguish between endogenous microglia or peripheral macrophages. Published data indicates an equivalent number of microglia and infiltrating peripheral macrophages in ischemic tissue at 72h post-HI, so given the fact that 80% of the Iba1+ cells are FtL+, this would likely indicate that both cell types can activate this response, but the relative contribution of phagocytic cells is not clear.^49,50^ Finally, with regards to iron sequestration response, the source of iron is not identified. While significant hemorrhage was not identified, peripheral iron could potentially increase. Published data has shown no increase to total brain tissue iron levels using this model, which suggests that the cellular changes in iron regulation are a reaction to dying cells spilling iron content into local milieu that requires sequestration to prevent ongoing oxidative damage.^17^

Contextualizing these findings with respect to the known differences between neonatal and adult stroke model also suggest that the neonatal brain is more susceptible to iron toxicity and lipid peroxidation during acute injury. Perls staining identifies iron deposition in the adult brain at 3-4 weeks after ischemic injury as opposed to the 4h post-HI observed in neonatal brains.^14,51^ During the first two post-natal weeks in rodents, cellular levels of labile iron, ferritin and iron import proteins are dynamically changing and with ferritin expression as well as labile iron decreasing over the first two post-natal weeks.^52–54^ These changes potentially make the neonatal brain more susceptible to iron toxicity as injury is occurring when the cells are most vulnerable to an acute increase in labile iron. Indeed, previous work has shown that ischemic injury in the neonatal brain generates more oxidative stress compared to the adult brain which has also been suggested to be due to developmental immaturity of anti-oxidant pathways.^55–57^ Over-expression of SOD1 which is able to neutralize oxygen free radicals by converting it into hydrogen peroxide is protective in adult stroke but exacerbates injury in neonatal mice.^58,59^ Thus, increased levels of peroxyl species, which can react with iron to create hydroxyl radicals, appear to be more toxic in the neonatal brain. Finally, oxidation of DHA specifically was recently identified as showing some selective vulnerability compared to arachidonic acid in an adult rodent stroke model, which is similar to that observed in this study.^60^ Lipid peroxidation appears to contribute to injury regardless of brain age, but there may be regional and age specific vulnerability to specific mechanisms that drive lipid peroxidation mediated cell death.

In conclusion, this study shows the temporal changes to cellular iron regulation and lipid peroxidation following neonatal-HI. The timing of these changes indicates that the toxic effects of iron and lipid peroxidation occur within the first 24h post-HI. These findings, indicate development of preventative strategies that prevent lipid peroxidation and iron accumulation may still be therapeutic options. Furthermore, augmenting or accelerating the iron sequestration response by microglia/macrophages or neuronal Gpx4 activity may be represent potential therapeutic strategies.

## Acknowledgements

We appreciate the services of the Vanderbilt Eicosanoid Laboratory for their analysis of isoprostanes and neuroprostanes.

Funding Sources: NIH/NEI K12EY015398 (JV), NIH/NINDS R01NS101156 (DMT), NIH/NEI R01EY028916 (JLD)

